# powerTCR: a model-based approach to comparative analysis of the clone size distribution of the T cell receptor repertoire

**DOI:** 10.1101/297119

**Authors:** Hillary Koch, Dmytro Starenki, Sara J. Cooper, Richard M. Myers, Qunhua Li

**Affiliations:** Department of Statistics, Pennsylvania State University, University Park, PA, USA; HudsonAlpha Institute for Biotechnology, Huntsville, AL, USA

## Abstract

Sequencing of the T cell receptor repertoire is a powerful tool for deeper study of immune response, but the unique structure of this type of data makes its meaningful quantification challenging. We introduce a new method, the Gamma-GPD spliced threshold model, to address this difficulty. This biologically interpretable model captures the distribution of the TCR repertoire, demonstrates stability across varying sequencing depths, and permits comparative analysis across any number of sampled individuals. We apply our method to several datasets and obtain insights regarding the differentiating features in the T cell receptor repertoire among sampled individuals across conditions. We have implemented our method in the open-source R package powerTCR.

**Author summary:** A more detailed understanding of the immune response can unlock critical information concerning diagnosis and treatment of disease. Here, in particular, we study T cells through T cell receptor sequencing, as T cells play a vital role in immune response. One important feature of T cell receptor sequencing data is the frequencies of each receptor in a given sample. These frequencies harbor global information about the landscape of the immune response. We introduce a flexible method that extracts this information by modeling the distribution of these frequencies, and show that it can be used to quantify differences in samples from individuals of different biological conditions.

## Introduction

Recent advances in high-throughput sequencing of the T cell receptor (TCR) repertoire provide a new, detailed characterization of the immune system. T cells, each displaying a unique TCR, are capable of responding to presented antigens and initiating an adaptive immune response. An immune response is described by rapid proliferation of T cell clonotypes whose TCRs are specific to the antigen. In humans, it is estimated that the body is capable of producing more than 10^18^ different TCRs [1, 2], where high diversity of the TCR repertoire implies a greater range of pathogens that can be fought off. A variety of studies have been published demonstrating the value in characterizing this immune response for purposes such as describing tumor cell origin [3] and predicting response to cancer therapy and infection [4]. The applications of TCR sequencing are many, but this type of data presents new needs for analysis techniques not met by existing tools for other kinds of genomic experiments.

Several groups have identified that the distribution of larger clone sizes in a sample can be approximated by a power law [5–8], which means that the number of clones of a given size decays approximately as a power of the clone size. This heavy-tailed distribution comes as a consequence of extensively proliferated clones actively participating in an ongoing immune response. More recent work has aimed to quantify statistically the diversity of the TCR repertoire, initially through the use of various estimators borrowed from ecology, such as species richness, Shannon entropy [9], and clonality. These estimators are known to be highly sensitive to sample size and missing observations. Given that the TCR repertoire is mostly populated by rare clonotypes, many of the clonotypes in the system are absent from any one sample. This presents a challenge to many of the ecological estimators. Model-based approaches to approximating the clone size distribution have also been proposed, with the goal of providing added stability and consequently more statistical power. Some examples are the Poisson-lognormal model [10], Poisson mixture models [11, 12], and a heuristic ensemble method [13]; however, these models lack a biologically meaningful interpretation, and further do not sufficiently account for the power law-like nature of the data. That is, power law distributions are heavier-tailed than the Poisson or even the lognormal distribution, leading to systematic bias in the model fit.

Previous research has also identified the imperfectness of the power law behavior for the clone size distribution below some clone size threshold [7, 8]. To handle the imperfectness, [7, 8] proposed to model large clones above the threshold using a type-I Pareto distribution, which is a member of the power-law distribution family, and omitted the clones with frequency below that threshold. The threshold is either user-specified or determined from the data based on a goodness-of-fit measure. Indeed, this model has certain biological basis. Through a stochastic differential equations setup that models the birth, death, selection, and antigen-recognition of cells active in the immune system, Desponds et al. [8] showed that the upper tail of the clone size distribution at equilibrium approximately follows a type-I Pareto distribution (Fig 1A). Unlike the Poisson and lognormal models, parameters in this model are related to relevant actors in the immune response, and can reveal certain biological insights into immune response, such as average T cell lifetime [8]. Yet, the resulting model excludes all clones below a certain frequency threshold.

**Fig 1.**
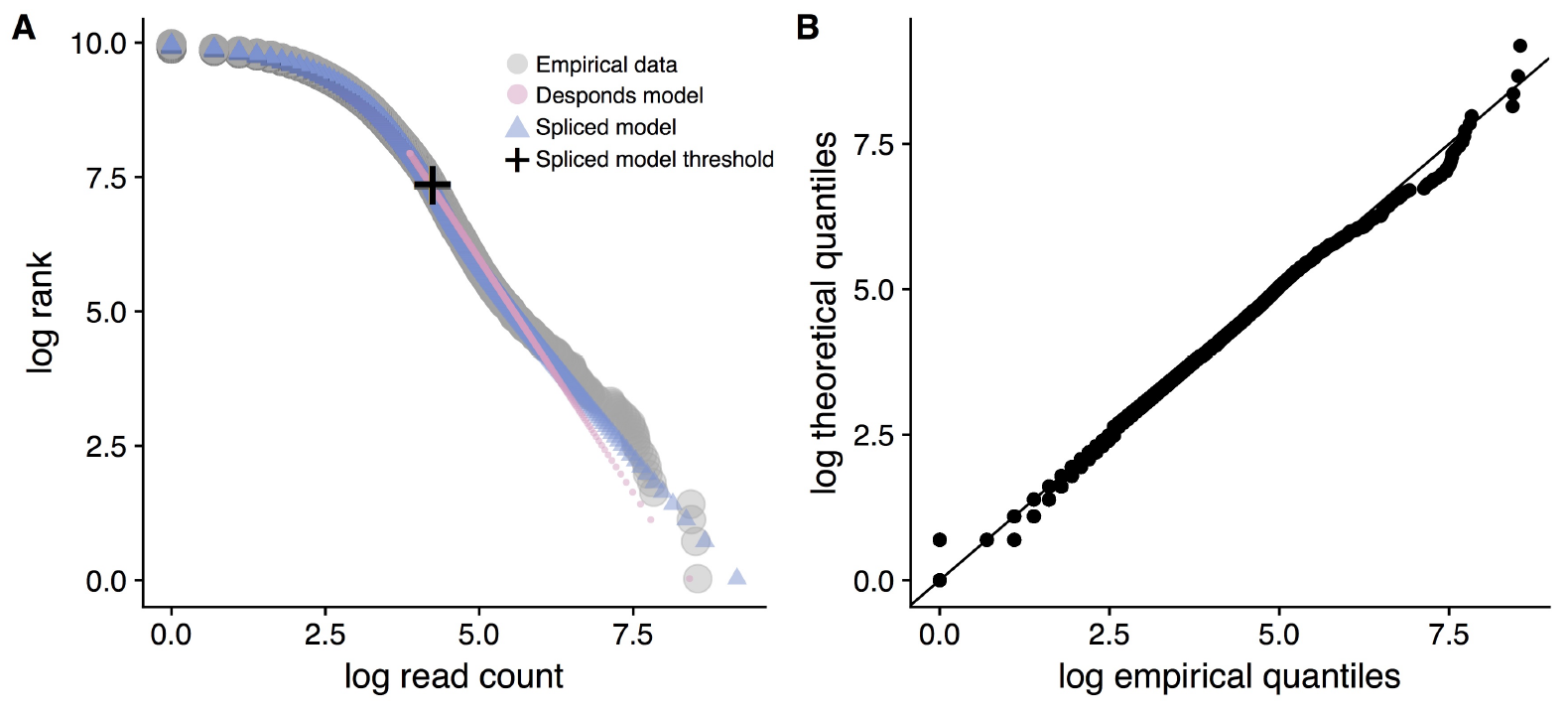
Examining the power law behavior of a sample clone size distribution. A: A sample repertoire (gray) of a Sarcoidosis patient and the fitted curves based on the Desponds method (pink) and our method (blue). The Desponds method only fits the data above the threshold it estimates (approximately 15% of the data). Our method fits all of the data. The cross marks the threshold estimated by our method. A complete collection of plots for every dataset analyzed are in S1. B: The QQ-plot of the theoretical fit using our model against the empirical data for the same dataset in A shows that our model can achieve a good fit.

However, even small clones may provide information; for example, Desponds et al. [8] indicated that the generation of new T cells affects the landscape of smaller clone sizes. Other studies have shown that low-frequency clones may support a diverse immune system and present a potential to mobilize against antigens, as in some cases having a clone size distribution highly dominated by a few clones has been correlated with unfavorable clinical outcome [14, 15]. With this in mind, we sought a means to exhaust all available data and consider modeling the complete clone size distribution.

To address this question, we propose a novel statistical tool, called powerTCR, to characterize the full distribution of the TCR repertoire. Our method models large clones that are above the threshold, where the power law begins, using the generalized Pareto distribution (GPD), which contains the type-I Pareto distribution as a special case, but provides a more flexible fit. It also models the small clones below the threshold using a truncated Gamma distribution. It determines the threshold in a data-driven manner simultaneously with the characterization of clone size distribution. Our final model contains parameters that are analogous to those found in the type-I Pareto model of Desponds et al. [8], relating our model to the biological interpretation of the dynamics of the immune system. Altogether, this allows our model to more accurately describe the shape of the clone size distribution for both large and small clones. Such a model is well suited for providing a global view of the state of the immune repertoire. It can also be employed to perform comparative analysis of healthy and compromised individuals to identify descriptors of strengths and deficiencies in the immune system.

## Results

### The discrete Gamma-GPD spliced threshold model

Our goal is to model the clone size distribution of a sample immune repertoire. Fig 1A shows a typical distribution plotted using the repertoire of a Sarcoidosis patient in [16]. If the data are truly Pareto distributed, this plot would appear linear [17, 18]. However, noting the linear behavior is only true for the far upper tail of the data, this suggests that these data are a departure from the Pareto distribution. This imperfect power law implicates the use of a heavy-tailed distribution above some threshold and a lighter-tailed distribution below that threshold. Here, we model the tail part with a GPD. The GPD, introduced by [19], is a classical distribution typically used to model the values in the upper tail of a dataset. This formulation results in a distribution with density

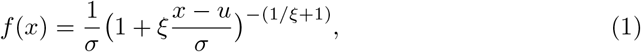

where *u* ∈ (–∞ *,∞*) is a threshold that typically needs to be prespecified, *σ* ∈ (0*,∞*) is a scale parameter, and *ξ* ∈(–∞*,∞*) is a shape parameter. The GPD has support *x ≥ u* when *ξ ≥* 0 and *u ≤ x ≤ u – σ/ξ* when *ξ <* 0. We model the bulk part with a Gamma distribution with the upper tail truncated at the threshold. The Gamma distribution has a flexible shape and can fit many different clone size distributions. The threshold and the parameters in the two distributions are estimated from the data simultaneously. This setup, where data above and below an unknown threshold are drawn from the “bulk” and “tail” distributions respectively, falls into a class of models called spliced threshold models. The typical motivation for the model is the belief that the data above and below the threshold are driven by different underlying processes. We refer the interested reader to [20] for a thorough review of the general spliced threshold model, and its applications in fields such as insurance, hydrology, and finance.

Denote the proportion of data above the threshold *u* as *φ*. Let the bulk model distribution function be *H_c_*(*x|θ_b_*) and the tail model distribution function be *G_c_*(*x|θ_t_*), where subscripts *b* and *t* denote the bulk and tail model parameter vectors, respectively. Then the distribution function of the model is given by

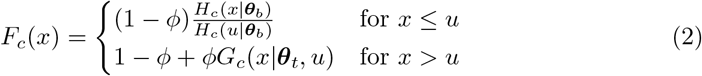

with corresponding density

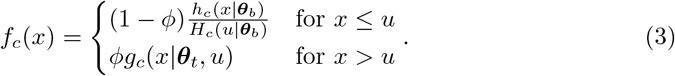

Because the clone size distribution is count data that typically exhibit numerous ties in the less frequently observed clonotypes, it is appropriate to treat this as a discrete problem. We modify the model in order to account for any quantized or censored data. Let *ψ* and Ψ be the density and distribution function of a continuous distribution, and let *d* be the interval length at which the data are censored. We obtain a quantized analog of *ψ* by letting

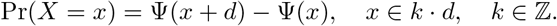

This results in a discrete model with distribution function

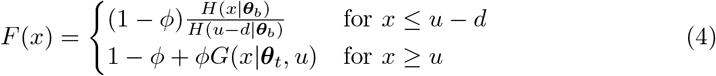

and corresponding probability mass function

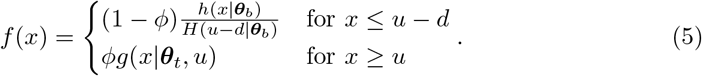

where *h*(*x|θ_b_*) ~discrete Gamma(*α, β*), *g*(*x|θ_t_*)~ discrete GPD(*u, σ, ξ*), and *d* = 1, which specifies that we model integer data (see Methods for the functional form of the discrete Gamma distribution and the discrete GPD). This discretization step turns out to be important for accurate estimation in our scenario. See S2 for a comparison between the performance of the discrete and continuous models in settings resembling true clone size distributions.

### Biological interpretation of the model

The relationship between the discrete Gamma-GPD spliced threshold model and the type-I Pareto model in Desponds et al. [8], hereafter referred to as the Desponds et al. model, allows us to draw connections between some of our parameters and the dynamics of immune response underpinning their approach. First, results from [8] show that the threshold at which the power law begins is indicative of the point over which a clone’s large size can be attributed to active immune response, as opposed to noise in the body that arises from processes such as self-recognition. The threshold fitted from the data provides an objective way to narrow down which clonotypes from a sample repertoire should be interrogated further. This notion is convenient for studying factors such as CDR3 (complementarity-determining region 3) amino acid motifs or specific V, D, and J genes important for combating certain antigens, which are typically determined based on a heuristic abundance cutoff. For example, [21] studies the 1,000 most abundant CDR3 amino acid motifs across all sampled peripheral blood mononuclear cell (PBMC) libraries, while [16] determines CDR3 amino acid motifs from clones that are present with 10 or more reads in a sampled repertoire. The threshold *u* estimated with our model, however, introduces a means to select motifs that does not rely on heuristics and automatically scales with sequencing depth.

Moreover, the shape parameter *ξ* of the GPD is inversely related to the shape parameter *α_d_* used in the Desponds et al. model (see Methods). As explained by Desponds et al., a small *α_d_*, i.e. a large *ξ*, implies increased average T cell lifetime and antigenic noise strength. They further show that antigenic noise strength grows as a consequence of a higher initial concentration of antigens and a higher rate at which new antigens are introduced. Interestingly, *ξ* also positively correlates with the familiar clonality estimator (1-Pielou’s evenness [22]). Indeed, as *ξ* increases, the clone size distribution becomes heavier-tailed—that is, more skewed towards dominating clones. This trend is in line with that of the clonality estimator, which favors a more uniform clone size distribution as clonality approaches 0 and a distribution dominated by expanded clones as clonality approaches 1. To numerically validate this relationship, we simulated the data from our model and computed the clonality (see Methods). We observed a high correlation between clonality and *ξ* (Spearman’s *ρ ≈* 0.9), confirming that *ξ* reflects the skewness towards dominating clones (see S3).

It is worth noting that our model acquires a theoretical gain via the threshold stability property of the GPD [23]. That is, for any generalized Pareto distributed data, the shape parameter *ξ* remains constant regardless of changes in *u*. In our context, this means that at decreasing sequencing depths, though the threshold *u* would decrease due to fewer cells being sampled, the shape parameter *ξ* in principle would be stable against the variation in sequencing depth. We will demonstrate this gain in stability on a murine tumor dataset. See Methods for our extension of the threshold stability property to the case of the discrete GPD.

In the following sections, we inspect three different datasets using our model. We compare our results to results from the Desponds et al. model to demonstrate the practical and theoretical benefits of our approach. We also make comparisons to results from the widely used richness, Shannon entropy, and clonality estimators. See Methods for information on computation of competing methods.

### Discrimination between tumors from MHC-II positive and control murine breast cancers

The expression of major histocompatability complex II (MHC-II) proteins in tumors correlates with boosted anti-tumor immunity. As part of a study of how MHC-II expression impacts tumor progression and functional plasticity of T cells [24], the CDR3 of TCR*β*-chains of tumor infiltrating lymphocytes (TILs) were sequenced from breast cancer tumor tissue from six BALB/c mice [25]. Three of the mice were grafted with MHC-II expressing tumor cells and three control animals received parental MHC-II-negative cells. Samples were collected at 21 days after the date of treatment.S4, Table 1 summarizes the number of unique clonotypes observed and the total number of reads in each sample.

### Our method quantifies response to treatment and differentiates treatment groups

We asked whether the MHC-II expression would cause a global change in the TCR repertoire of the host mice. We analyzed the TCR repertoire in the individuals’ tumor tissue using our proposed model and the Desponds et al. model. S4, Tables 2 and 3 show parameter estimates from our model and the Desponds et al. model, respectively. Results from our model support a claim that increased expression of MHC-II induces an increased rate of clonal expansion at the tumor site, as indicated by the uniformly larger estimates of the tail shape parameter *ξ* in the treatment group.

Because the TCR repertoire is often used as an indicator of an individual’s immune potency, we next evaluated how well our method discriminates between samples from case and control groups. We compared our method to the Desponds et al. model and the richness, Shannon entropy, and clonality estimators. For our model, we computed the pairwise Jensen-Shannon distance (JSD) between the fitted distributions of each pair of samples, and then used it as the distance measure to cluster samples with hierarchical clustering (see Methods) using Ward’s method. Clustering of the Desponds et al. model was done similarly by using a generalization of JSD to continuous distributions, trading the summation for an integral. For all ecological measures, we computed pairwise Euclidean distances between estimates which we then used for hierarchical clustering of the samples. Fig 2 shows that our approach clusters the data by treatment and control groups. This is contrasted against all other methods, which display an incorrect construction of the expected true relationships among the experimental subjects. In the case of the ecological estimators, poor clustering likely occurs because a one-number summary of repertoire diversity cannot capture intricacies in the data, and varying sequencing depths across samples may further bias results. In the case of the Desponds et al. model, on the other hand, poor clustering is likely a result of the lack of a robust fitting procedure and a less flexible model. Moreover, the exclusion of smaller clones reduces power to discriminate between samples. See S5 for a demonstration of reduced clustering performance when only using the tail part of our spliced model.

**Fig 2.**
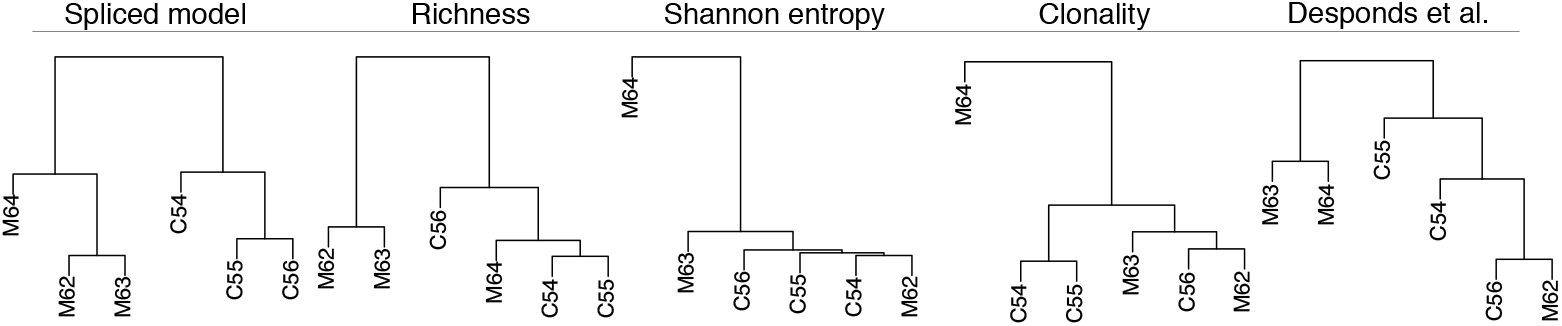
Hierarchical clustering of mouse tumor samples based on different quantifications of the TCR repertoire. The MHC-II-positive individuals are labeled with an ‘M’, while the control individuals are labeled with a ‘C’.

### Our results are robust to variation in sequencing depth

One challenge with TCR experiments is that the sequencing depth directly affects the number of TCR variants that are discovered. It is not simple to obtain the same sequencing depth across individuals, and it is also undesirable to discard data in favor of maintaining an equal number of reads across TCR repertoire libraries. An ideal scheme for comparative analysis achieves stable classification across different sequencing depths.Thus, we compared the stability of our model against the Desponds et al. model. To do this, we randomly downsampled reads in the mouse data to 80, 60, 40, and 20% of total reads from the original samples. We fit our model and the Desponds et al. model in each case, then performed hierarchical clustering according to JSD. The clustering induced using our model was the same at every downsampling level, while clustering induced using the Desponds et al. model changed each time. S6 contains the dendrograms from both models at each downsample level, but Fig 3 summarizes this information by illustrating the relative JSD between sample C54 and the remaining samples across downsample levels. The spliced threshold model clearly maintains a similar trend across downsample levels, not to mention that the relationship inferred distinguishes between treatment and control groups. However, the downsampling study reveals that relative distances between sampled individuals observed using the Desponds et al. model are quite erratic.

**Fig 3.**
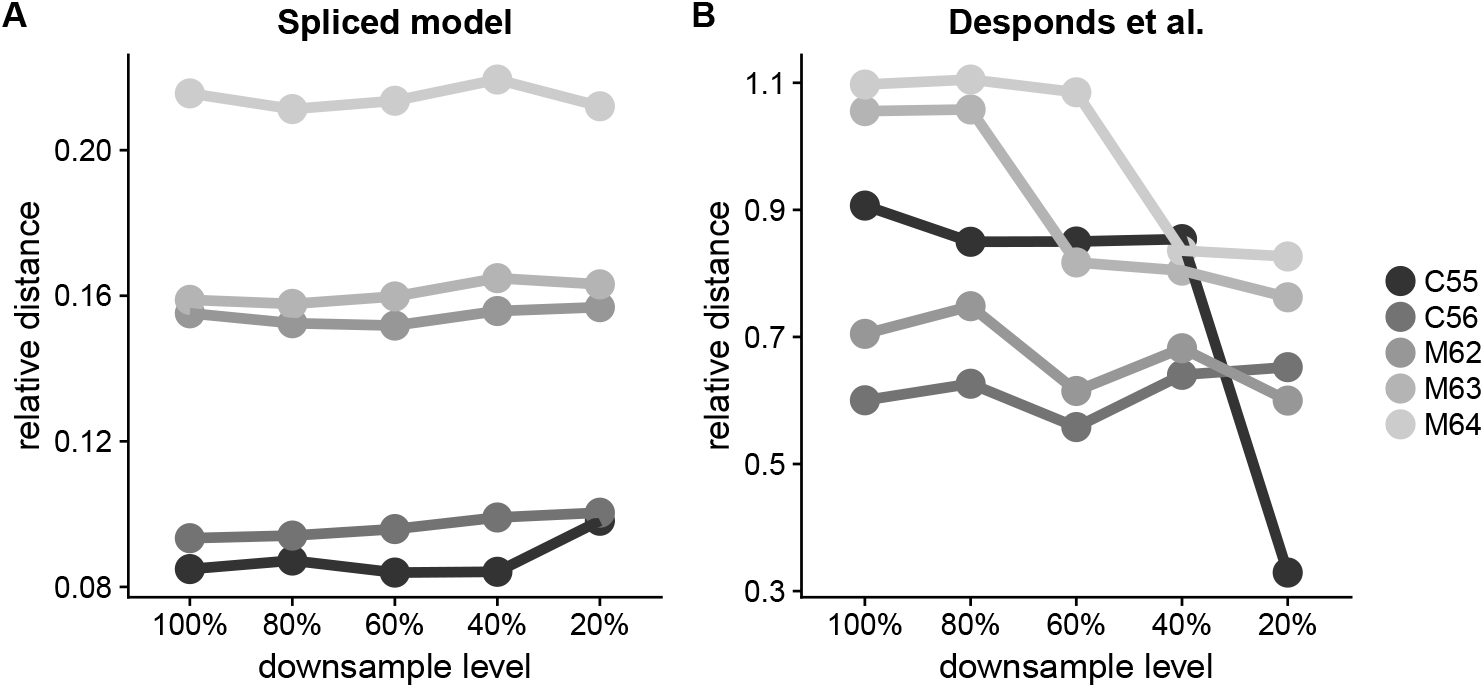
Robustness to variation in sequencing depth on the mouse tumor data. Relative JSD between sample C54 and the remaining samples across downsampling levels, using A: our model and B: the Desponds et al. model.

The higher stability of our results is due in part to the threshold stability property of the GPD. It is also attributed to the fact that our method models the full clone size distribution. Doing so circumvents the threshold selection problem encountered when applying the Desponds et al. model, a recurrent issue when fitting models such as the type-I Pareto distribution [26]. For their approach, Desponds et al. select the threshold that minimizes the Kolmogorov-Smirnov statistic [27], a commonly used goodness-of-fit strategy for fitting a power law distribution [28]. However, the results on this dataset show that this strategy does not always yield stable results across different sequencing depths.

### Differentiation between subtypes of Sarcoidosis patients

Sarcoidosis is an inflammatory disease that typically is accompanied by an accumulation of activated CD4+ T cells in the lungs. A particularly acute form of Sarcoidosis, called Löfgren’s syndrom (LS), occurs with additional, more severe symptoms. A known signature of LS is the bombardment of the lungs with CD4+ T cells, which is expected to significantly alter the entire landscape of the TCR repertoire. We applied our method to TCR repertoire data of LS and non-LS Sarcoidosis patients [29], originally described in [16]. In this study, bronchoscopy with the bronchoalveolar lavage was performed on a cohort of 9 LS and 4 non-LS individuals and prepared for TCR *α* and *β* chain sequencing.

We compared the TCR distribution between LS and non-LS Sarcoidosis patients using our method and the competing methods. In order to visualize closeness of samples, we generated a distance matrix using JSD between fitted distributions using our method and the Desponds et al. model. The estimated parameters are in S4, Tables 2 5 and 6, respectively. We then applied non-metric multidimensional scaling (MDS) to 2 the distance matrix and plotted the first two coordinates. For the ecological estimators,2 we simply plotted centered and scaled estimates. As shown in Fig 4A, results from our 2 model cluster LS patients into a tight group distinct from non-LS patients, bolstering 2 the claim that LS patients exhibit a signature immune response. On the other hand, competing methods fail to uncover any pattern (Fig 4B–C).

**Fig 4.**
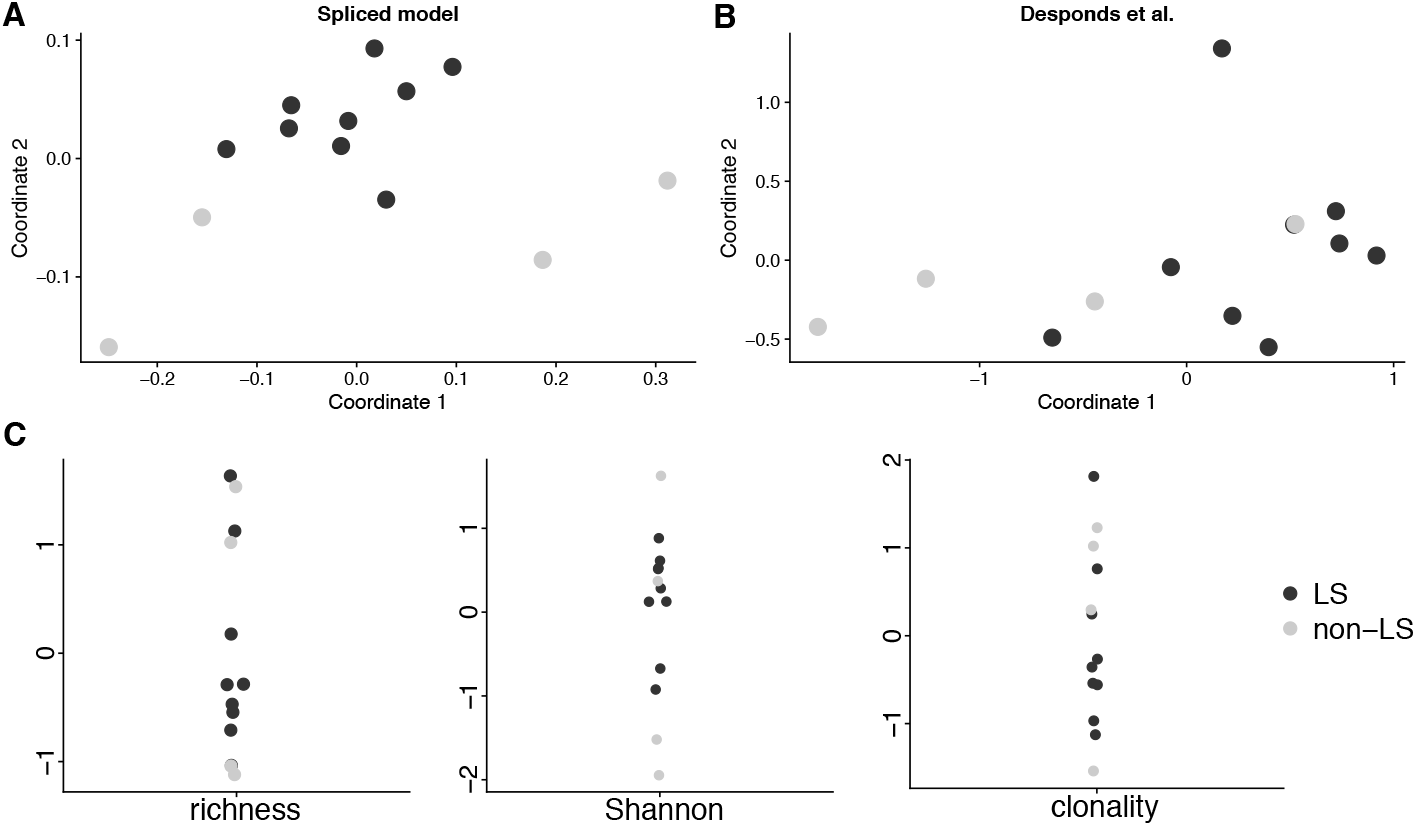
Differentiation of TCR repertoires between LS and non-LS Sarcoidosis patients. A–B: Multidimensional scaling representation of distances between TCR repertoires computed with JSD for the spliced threshold model and the Desponds et al. model. C: Centered and scaled estimates of richness, Shannon entropy, and clonality.

### Relationship between the landscape of the clone size distribution and clinical outcome in glioblastoma patients

We applied our method to data collected during a clinical trial of 13 glioblastoma patients receiving autologous tumor lysate-pulsed dendritic cell (DC) vaccine therapy [30], first detailed in [31]. Three intradermal injections were administered to patients at biweekly intervals. TCR*β*-chains from PBMC samples were sequenced for the patients prior to vaccinations and two weeks following the final injection. Patients were followed up with and their time to progression (TTP) and overall survival (OS) were recorded. TTP was defined as the time from the first DC vaccination until MRI-confirmed tumor progression. OS was calculated as the time from the first DC vaccination until the patient’s death from any cause. We investigated whether current tools using TCRs sequenced only from blood samples indicate anything about patients’ survival time and time to progression.

We first fit our model to the pre- and post-treatment samples. In both cases, we classified the patients into two groups using the hierarchical clustering based on our model, the Desponds et al. model, and the richness, Shannon entropy, and clonality estimators. No clear grouping with respect to either TTP or OS could be observed from any clustering on the pre-treatment samples, whether by the model-based methods or the selected estimators (see S7). However, among post treatment samples, our method tends to cluster together patients with better clinical outcome (Fig 5A). This may indicate that the DC therapy alters the landscape of the TCR repertoire into a form that promotes favorable clinical outcome.

**Fig 5.**
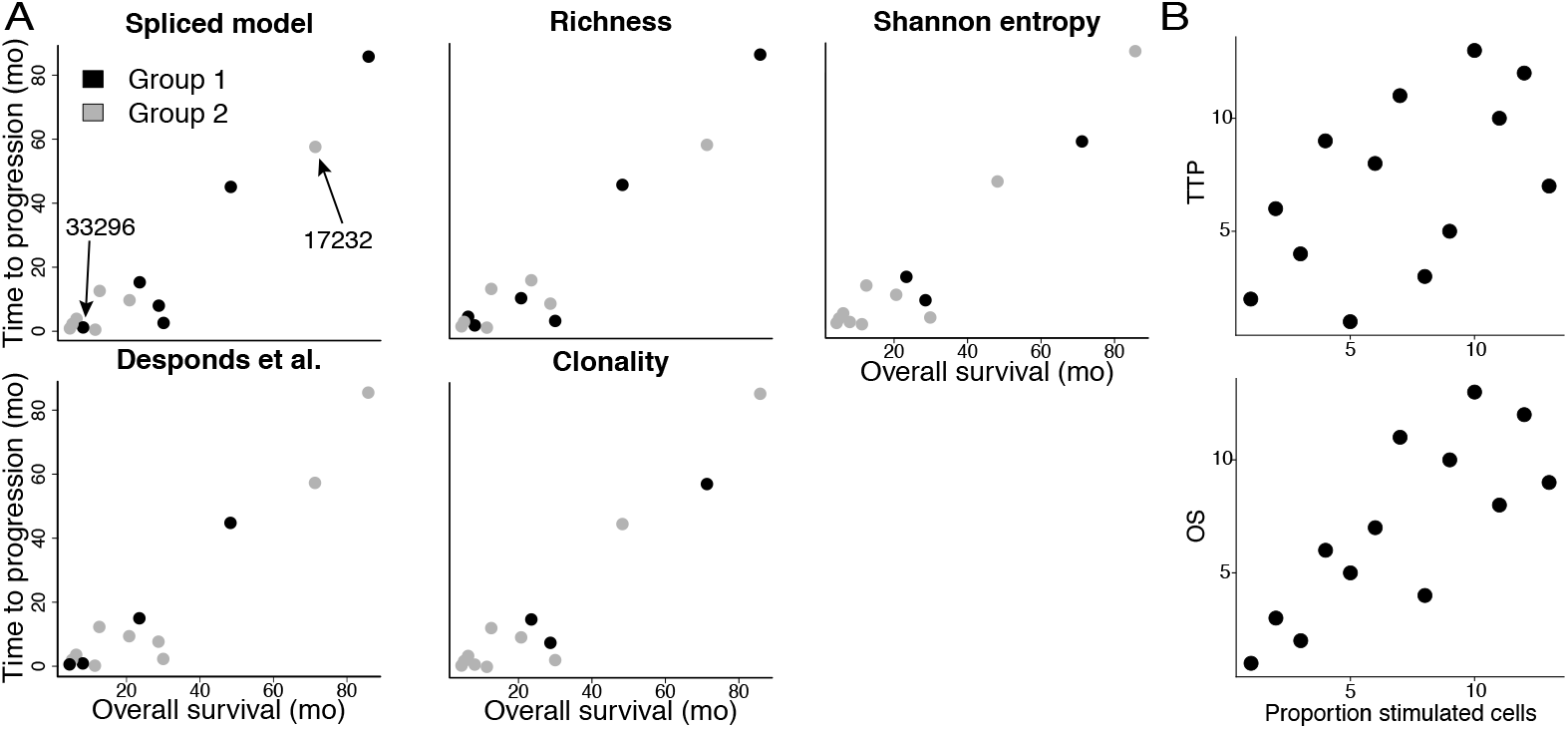
Association with time to progression (TTP) and overall survival (OS) time in glioblastoma patients. A: Patient classification based on TCR repertoire quantification and its relationship to TTP and OS. Patients are classified into two groups (black and gray) using hierarchical clustering according to the estimates of TCR repertoires from the post-treatment samples. Anomalous patients 33296 and 17232 are called out on the Spliced model plot. B: Relationship between the proportion of highly stimulated clones inferred by our method and TTP or OS.

We do, however, cluster one patient (ID: 33296) with low TTP and OS in the group with overall higher TTP and OS. Interestingly, this misplaced patient had the lowest estimated TIL count and tumor/PBMC overlap of the entire cohort (S4, Table 7). Tumor/PBMC overlap was defined as the total number of reads of shared CDR3s normalized by total reads in the tumor and PBMC samples. Similarly, patient 17232 displayed among the best clinical outcome but clustered with lower-performing patients. Patient 17232 had the highest TIL count and level of tumor/PBMC overlap in the whole cohort. This information taken as a whole suggests that, while the clone size distribution found in blood may indicate something about a patient’s response to treatment, it still does not guarantee that T cells will infiltrate the tumor, an important factor for clinical benefit [32]. S8 highlights the clone size distributions of these two patients against all others.

Notably, inferred thresholds (minimum *u* = 4, maximum *u* = 6) on this dataset are much lower than on other datasets. This is likely because this dataset contains less deeply-sequenced samples than the others, which consequently reduces the threshold.

Noting that clones with size at or above the estimated threshold are considered active participators in the immune response, we sought to investigate whether any relationship existed between clinical outcome and the proportion of more highly stimulated cells. We defined the proportion of highly stimulated cells to be the total number of reads at or above the threshold, normalized by the total number of reads in the entire repertoire (S3, Table 9). We found correlations between this measure and both TTP (Spearman’s *ρ* = 0.54) and OS (Spearman’s *ρ* = 0.80). Rank scatterplots for these correlations are in Fig 5B. The positive correlation we uncovered suggests that this statistic could be a useful tool to quantify the antigen-specificity of the sample.

### Relationships among sorted CD4^+^ T cell subtypes in individuals with type 1 diabetes and healthy donors

Risk factors for type 1 diabetes (T1D) are known to be heritable, yet genes alone are not sufficient explanation for drivers of the disease. Studies of monozygotic twins have revealed that, given one twin has T1D, the other will only have it at most half of the time [33]. The CD4^+^ T cell is viewed as the initiator of T1D as dysregulation of CD4^+^ antigen-recognition drives the autoimmune disease. Seeking out apparently non-heritable determinants of T1D, [34] conducted a deeper investigation of the CD4^+^ T cell. Briefly, the authors obtained PBMCs from 14 volunteer healthy donors (HDs) and 14 recently diagnosed patients with T1D. The cells were sorted using flow cytometry into distinct T cell subsets (true naїve; TN, central memory, CM; regulatory, Treg; and stem cell-like memory, Tscm) and TCR*β*-chains were sequenced. The authors conducted a thorough analysis, finding shorter CDR3 sequence lengths and lower overall repertoire diversity among patients with T1D. However, on a per-individual basis, the authors were unable to uncover a relationship between repertoire diversity and disease status. Since the the spliced threshold model provides a new means to probe this complex data, we applied our approach to complement the original analyses.

#### Trajectory in clone size distribution across CD4^+^ T cell subtypes

While [34] considered pairwise correlations in diversity indices of different CD4^+^ T cell subsets from the same individual, their analysis provides no visualization of the data as a whole. Our method for comparative analysis, combined with MDS, allows this to be done naturally. Like the Sarcoidosis patient analysis, we applied our method and competing methods, and visualized the results as before (Fig 6A–C).

**Fig 6.**
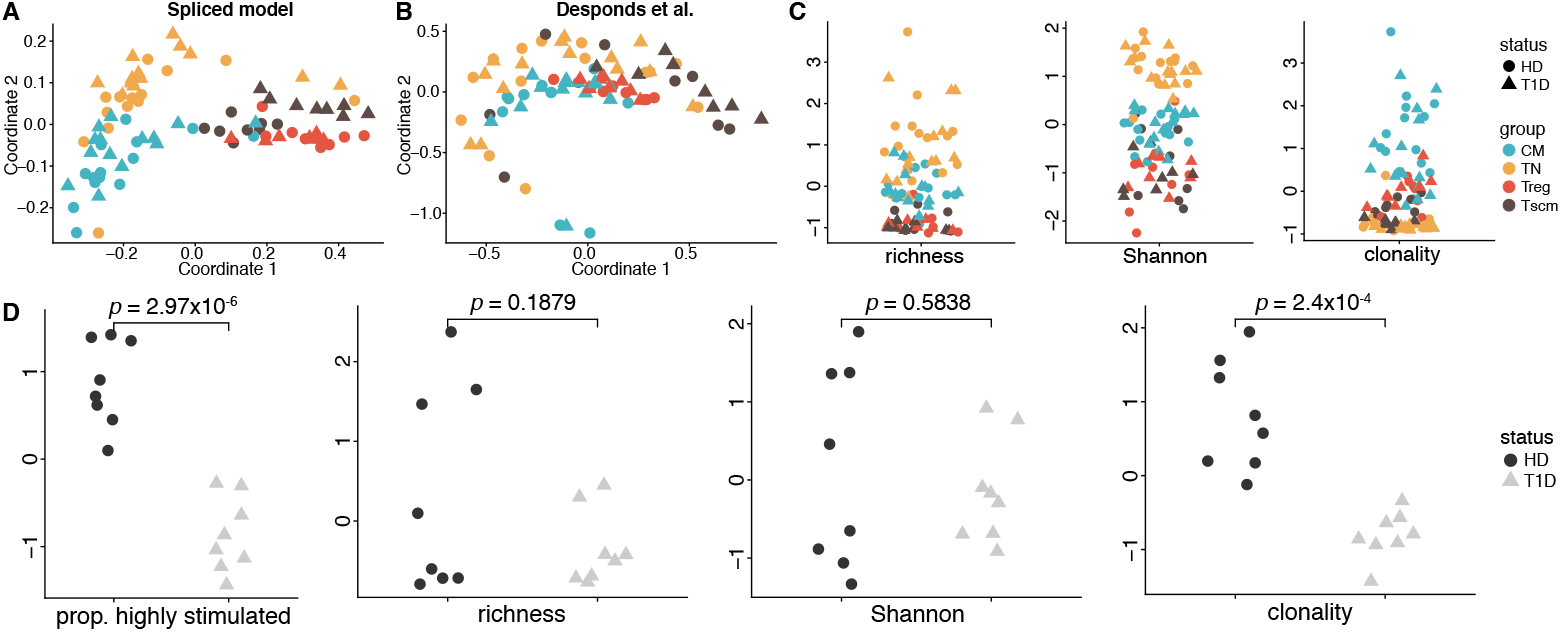
Trajectory of TCR repertoires among CD4^+^ T cell subsets between T1D patients and HDs. A–B: MDS representations of JSD between TCR repertoires from the spliced threshold model and Desponds et al. model. C: Centered and scaled estimates of richness, Shannon entropy, and clonality for all CD4^+^ T cell subsets. D: The proportion of highly stimulated clones derived from our method and centered and scaled diversity estimates in the Tscm subset.

It is known that TN cells propagate into both Tscm and CM cells, Tscm cells also propagate in CM cells, and Treg cells generally originate separately in the thymus [35–37]. Our approach appears to support a gradual change in repertoire shape throughout this differentiation process, as a clear trajectory is revealed in the MDS representation of the JSD between fitted models in Fig 6A. On the other hand, the MDS representation obtained using the Desponds et al. model does not separate results by cell type or donor status. The ecological estimators recapitulate some results found in the original analysis [34], for example that TN are more diverse than other cells, but 3 uncover nothing more.

Upon closer examination of the results from our model, we noticed that individuals from the T1D and HD groups are not well-separated in the cell types in Fig 6A, except in the Tscm subset. Tscm cells, identified *in vitro* in [35], are a stable, yet multipotent, subset of cells sustained via proliferation and turnover throughout the human lifetime and have been suggested to play a major role in establishing memory in immune response [35, 36]. More recently, Tscm cells have been implicated as a factor in development and treatment of autoimmune disorders such as acquired aplastic anemia [38] and systemic lupus erythematosis [39]. This motivated a deeper evaluation of the Tscm subset. Recalling that the proportion of highly stimulated clones, an estimator derived from our model, proved informative in our analysis of the TCR repertoires sequenced from PBMCs of individuals with glioblastoma, we decided to again apply it here to the Tscm subset alongside the ecological estimators.

As shown in Fig 6D, richness and Shannon entropy have difficulty differentiating the T1D and HD groups (*p* = 0.1879 and *p* = 0.5838 for a two-sided *t*-test, respectively), yet clonality and our measure uncover a clear split between the two groups (*p* = 2.4 × 10^*−*4^ and 2.97 × 10^*−*6^, respectively). This indicates the potential for the Tscm TCR repertoire to be used as a biomarker for detecting the status of T1D, suggesting the relevance of Tscm cells in T1D pathogenesis.

## Discussion

We have developed a model, the discrete Gamma-GPD spliced threshold model, and demonstrated its utility on several datasets. As shown in our analyses, several biologically relevant descriptive features can be obtained from our model. One is the tail shape parameter *ξ*, a measure of the weight of the upper tail of the clone size distribution, where a heavier tail of the fitted model implies a more dominated distribution of expanded clones. Another is the proportion of total reads at or above the estimated threshold, a possible measure of intensity of the immune response. The third is the estimated threshold, which is a useful guide to objectively identify CDR3 motifs for downstream analysis. This could involve denoting motifs as only those CDR3s found in TCRs with frequencies at or above the estimated threshold for a given sample, or it could mean studying TCR gene usage among that same group of clonotypes. Though the dynamics driving our model form a compelling argument for this interpretation of the threshold, we acknowledge that further biological validation on more datasets is still needed to confirm this.

Similar to other estimators, our model requires that a repertoire be adequately sampled. Without adequate sampling, the differentiating features between TCR repertoires will be masked [8], and the estimated model parameters will not be reliable. Given the immense diversity of the TCR repertoire, one should in general be cautious about using any method to make inference about a sample TCR repertoire when few cells are sequenced. With sufficient samples, though, the spliced threshold model provides the user a meaningful high-level view of the TCR repertoire.

The diversity of the TCR repertoire and its responsiveness to stimuli provide a high-dimensional biomarker for monitoring the immune system and its adaptivity. Robust assessment of the clone size distribution through TCR sequencing is important for understanding this diversity. The discrete Gamma-GPD spliced threshold model is a flexible model that effectively captures the shape of the clone size distribution. It is especially appropriate since the heavy-tailed GPD is a good fit to model the highly expanded clones that dominate many TCR repertoire samples. The method also provides a means to comparatively analyze a collection of TCR repertoire samples while maintaining convenient theoretical properties and interpretations.

Compared with existing approaches, our method is more flexible, utilizes the full clone size distribution, is less sensitive to sequencing depth, and identifies the threshold in a data-driven manner. The parameters estimated from our method are biologically relevant and instructive to the dynamics of immune response. Our results on multiple datasets also show that the spliced threshold model is powerful in a range of scenarios for comparing TCR repertoires across samples, revealing potential trends in the landscapes of clone size distributions of affected immune systems.

## Methods

### Estimation

We use maximum likelihood to estimate the parameters of our model. First, we more explicitly specify the form of our distribution. Letting *x~* Gamma(*α, β*), we write the probability mass function of a discrete Gamma distribution as

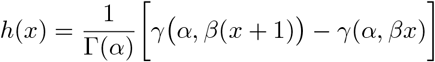

for *α* > 0, *β* > 0, ∊ ℤ, and where *γ*(*α, βx*) is the lower incomplete gamma function

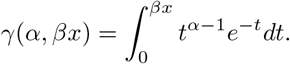

If *x~* GPD(*u, σ, ξ*), we write the probability mass function of a discrete GPD as

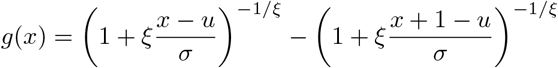

for *u* (–∞*,∞*), *σ* ∈ (0*,∞*), and *ξ* ∊ (–∞*,∞*). The discrete GPD has support *x ≥ u* when *ξ ≥* 0 and *u ≤ x ≤ u – σ/ξ* when *ξ <* 0, where *x* ∊ ℤ. In all analyses presented here, we make no assumptions on the sign of *ξ*, although empirically we tend to observe *ξ* > 0.

To proceed, we employ a profile likelihood approach. Let *u* be the threshold, *θ_b_* be the bulk parameter vector *{α, β}*, *θ_t_* be the tail parameter vector *{σ, ξ}*, and *θ* be the parameter vector {*θ_b_, θ_t_*}. Let also *h* and *H* be the density and distribution function of a discrete Gamma distribution, respectively, and let *g* be the density of a discrete GPD. Then the complete data likelihood is given by

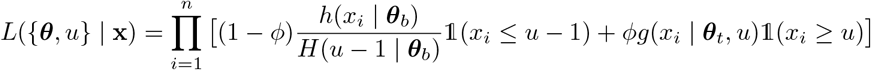

and the profile likelihood of the model at *u* is denoted as

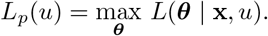

A grid search over a suitable range of thresholds u*^*^* = (*u*_1_*, …, u_k_*) may be implemented to maximize the profile likelihood. In this study, we adopted an approach similar to those of [19] and [40], searching for thresholds at or above the 75% quantile of the sample. The estimated parameters are then

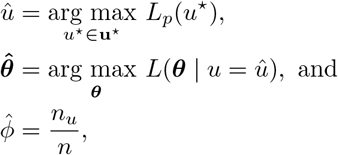

where *n* is the total number of clones and *n_u_* denotes the number of clones with size greater than or equal to the threshold.

### Computation for competing methods

The Desponds et al. model was fit as previously described [8]. Briefly, the model has density

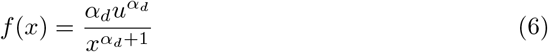

and distribution function

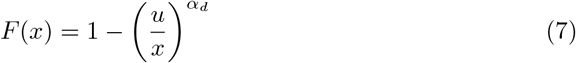

where *u* > 0 is the threshold and *α_d_* > 0 is a shape parameter. For each sample TCR repertoire, a grid of potential thresholds u^*^ = (*u*_1_*, …, u_k_*) was constructed by considering every unique clone size in the repertoire. Then, for each *u_i_*, the shape parameter is estimated as

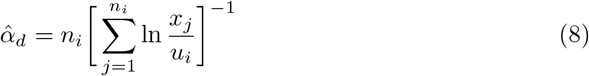

where *n_i_* is the number of clones with size larger than the threshold *u_i_*. Once this value is computed for every threshold in **u^*^**, the threshold and corresponding *α̂* were chosen to minimize the Kolmogorov-Smirnov statistic.

The ecological estimators [9, 22] were computed as follows. For a sample *X*, let *S*(*X*) be the sample richness, defined as the number of unique clonotypes in *X*, and let *p_i_* be the number of cells of clonotype *i* normalized by the total number of cells in the sample.Then, the Shannon entropy of *X* is

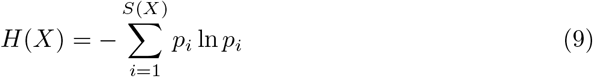

and the clonality of *X* is

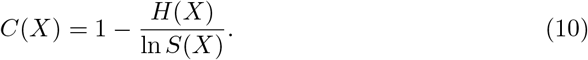

### *ξ* is inversely related to the the shape parameter of the type-I Pareto distribution

The Desponds et al. model, which is a type-I Pareto distribution, and the “tail” part of our model, which is a GPD, are closely related. In fact, the GPD contains the type-I Pareto distribution as a special case. We can write the distribution function of the *y*, where *y ~* GPD(*u, σ, ξ*), as

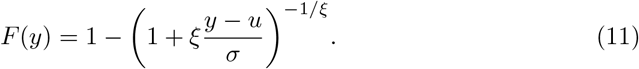

Now, let 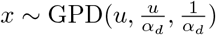. Then

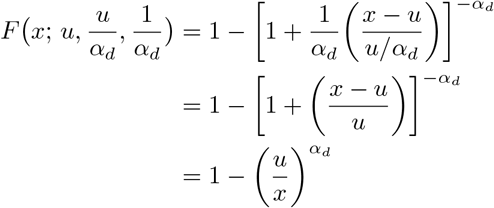

which is exactly the distribution function of a type-I Pareto distribution with threshold *u* and shape *α_d_* (Eq 7). Of course, this exact relationship only holds when 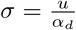. Nevertheless, *α_d_* and *ξ* perform the same function in their respective distributions, adjusting the weight of the tail. This relationship always holds – a larger *ξ* (smaller *α_d_*) implies a heavier-tailed distribution, while a smaller *ξ* (larger *α_d_*) implies a lighter-tailed distribution.

### Correlation between *ξ* and clonality

We conjecture that *ξ*, the shape parameter of the GPD, positively correlates with clonality. We numerically validated this claim using a simulated cohort of 48 clone size distributions. That is, we generated samples of *n* = 20, 000 clonotypes, where our 48 parameter settings were derived from every combination of *α* ∊ {3, 5, 10}, *ξ* ∊ {.25, .5, .75, 1.1}, and *φ* ∊ {0.1, 0.15, 0.2, 0.25}. We chose *β* = 0.15, 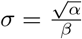, and *u* = *Q_α,β_*(1 – *φ*) in each simulation, where *Q_α,β_* is the quantile function of the Gamma distribution with mean 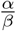. To adjust for the effect of sample size on clonality, we downsampled the simulated data so that each sample contained the same number of reads (415,989 total reads per sample). We computed the clonality of each simulated TCR repertoire on these adjusted datasets.

### Comparative analysis of multiple samples using the spliced threshold model

The relationship between a pair of TCR repertoires can be elucidated by evaluating the distance between their fitted spliced threshold models. Several methods to compare densities are available. We propose measuring the distance between each pair of distributions using Jensen-Shannon distance (JSD) [41]. This metric is a symmetric and smoothed adaptation of the well-known Kullback-Leibler divergence that does not require the distributions under comparison to share the same support.

Given discrete distributions *P* and *Q*, the JSD between *P* and *Q* is

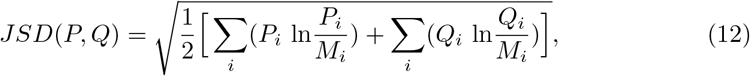

where 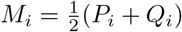. The resulting distances allow analysis and visualization via 4 MDS or hierarchical clustering of the samples. Throughout our study, we use Ward’s method for hierarchical clustering.

### Threshold stability of the discrete GPD

The threshold stability property of the GPD is well-established [23]. Here, we show that the property also holds for the discrete GPD. Let *X ~* discrete GPD(*u, σ, ξ*) and denote its distribution function as *F* with *F_c_* as its continuous analog. Then we can write

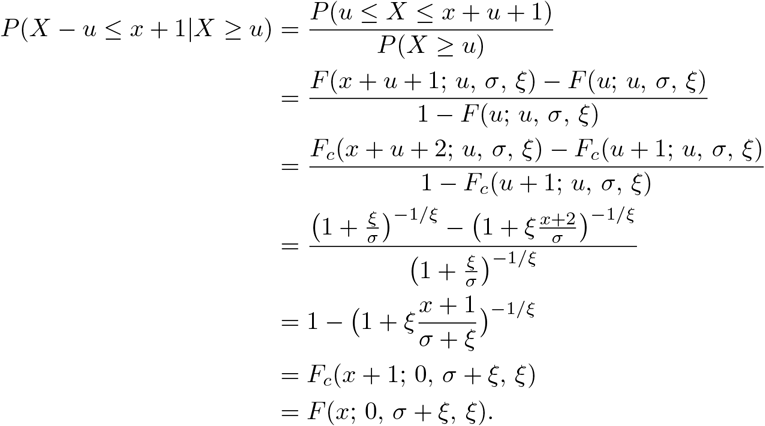

This states that if *X* discrete GPD(*u, σ, ξ*), then *X – u ~* discrete GPD(0*, σ* + *ξ, ξ*). Or, for our application, consider a clone size distribution, where clones larger than some threshold *u* are distributed according to the discrete GPD. At decreasing sequencing depths, this estimated *u* decreases, implying naturally that the size a clone in the sample must achieve to be considered “expanded” decreases. Still, while *u* shrinks, the threshold stability property states that *ξ* remains constant.

## Supporting information

**S1. Visualization of all model fits.** For each real data sample, a plot analagous to Fig 1A is provided.

**S2. Simulation for comparing the accuracy of parameter estimation for the discrete spliced threshold model, the Desponds et al. model, and the continuous spliced threshold model.** Details on simulation study comparing the discrete model and Desponds et al. model, and simulation study comparing the discrete 4 and continuous models.

**S3. Correlation between *ξ* and clonality.** Information on simulation study that finds strong positive relationship between *ξ* and the clonality estimator.

**S4. Summaries for the four TCR repertoire datasets.** Tables summarizing number of unique clonotypes per sample, total reads sequenced per sample, and other patient-specific information for the Sarcoidosis and glioblastoma datasets. Additionally, tables containing fitted model parameter estimates for both our model and the competing model, as well as ecological estimator values, computed for every sample.

**S5. Comparative analysis using the GPD.** By comparing results from our full model to those from only our tail model, we observe empirically the gains from including the full clone size distribution.

**S6. Dendrograms for downsampling study.** We downsampled mouse tumor data to 80, 60, 40, and 20% of total reads. We used JSD to compute pairwise distances between the samples for our model fits and the Desponds et al. model fits at each downsample level and did hierarchical clustering using Ward’s method. The dendrograms for each model at each downsample level are presented here.

**S7. Clustering of pre- and post-treatment glioblastoma patient samples.** Clustering dendrograms generated on pre- and post-treatment glioblastoma samples. Groupings presented for the post-treatment samples here correspond to the colored groupings in Fig 5.

**S8. Clone size distributions of glioblastoma patients.** This plot calls out patients 33296 and 17232 from the glioblastoma patients. Patient 33296 incorrectly clustered with the individuals with favorable clinical outcome, while patient 17232 incorrectly clustered with individuals with unfavorable clinical outcome.

## Data Availability

The murine breast cancer data and Sarcoidosis patient data analyzed in this study are available at Gene Expression Omnibus (http://www.ncbi.nlm.nih.gov/geo) under accession numbers GSE119670 and GSE100378, respectively. The glioblastoma and T1D data analyzed in this study are available in the immunoSEQ Analyzer®database (https://clients.adaptivebiotech.com). Our method is implemented in the R package powerTCR, freely available on BioConductor (https://bioconductor.org/packages/devel/bioc/html/powerTCR.html) under an Artistic-2.0 license.

